# Antibody-dependent DENV infection induces distinct transcriptomic responses in infected and bystander human macrophages

**DOI:** 10.1101/2024.07.11.603036

**Authors:** Céline S. C. Hardy, Adam D. Wegman, Mitchell J. Waldran, Adam T. Waickman

## Abstract

Dengue virus (DENV) is a mosquito-borne flavivirus which coexists as four genetically and immunologically distinct serotypes (DENV1-4). In secondary heterologous DENV infection, pre-existing immunity is believed to contribute to severe disease through antibody dependent enhancement (ADE). Although the elevated pathology observed in ADE conditions has been described, the cell intrinsic mechanisms governing this process remain unclear. Using scRNAseq, we investigated the transcriptomic profiles of human monocyte-derived macrophages infected by DENV via ADE compared to conventional infection conditions. Unsupervised analysis of scRNAseq data enabled the identification and differentiation of infected and bystander/uninfected cells in a heterogeneous cell culture. Differential gene expression and Ingenuity Pathway analyses revealed a number of significantly up- and down-regulated genes and gene networks between cells infected by ADE compared to conventional infection. Specifically, these pathways indicated mechanisms such as suppressed interferon signaling and inflammatory chemokine transcription in cells infected via ADE. Further analysis revealed that transcriptomic changes were independent of viral RNA within infected cells, suggesting that the observed changes are reflective of cell-intrinsic responses and not simply a function of per-cell viral burden. Bystander cells in ADE conditions also demonstrated distinct profiles, indicating an immunologically activated phenotype enriched for the expression of gene networks involved with protein translation, cytokine production, and antigen presentation. Together, these findings support the concept that DENV infection via ADE induces a qualitatively different transcriptomic response in infected and bystander cells, contributing to our understanding of ADE as a mechanistic driver of disease and pathogenesis.

**IMPORTANCE:** Dengue virus (DENV) is a mosquito-borne human pathogen with a significant and growing global health burden. Although correlates of severe dengue disease are poorly understood, pre-existing immunity to DENV has been associated with severe disease risk and known to contribute to an alternative route of viral entry termed antibody-dependent enhancement (ADE). Using single cell RNA sequencing, we identified distinct transcriptomic processes involved in antibody-mediated DENV entry compared to conventional receptor mediated entry. These data provide meaningful insight into the discrete processes contributing to DENV pathogenesis in ADE conditions.

## INTRODUCTION

Dengue virus (DENV) is a mosquito-borne flavivirus and globally relevant human pathogen accounting for an estimated 390 million infections yearly^1,2^. DENV exists as four genetically and immunologically distinct serotypes (DENV1-4) which co-circulate in endemic areas, primarily in tropical and subtropical regions. Clinical presentations of dengue range from an acute febrile illness to life-threatening hemorrhagic fever characterized by plasma leakage, hemorrhage, and severe shock^1,3,4^. Prognostic biomarkers of severe dengue are poorly defined, although the best described risk factor for severe disease is a secondary DENV infection with a heterologous serotype^5,6^. One of the mechanisms believed to contribute to the increased risk of severe disease in secondary heterologous infection is antibody-dependent enhancement (ADE), a process by which pre-existing antibodies facilitate viral entry into phagocytic leukocytes by Fc gamma receptor (FcγR)-mediated endocytosis^7–13^. The presence of discrete pathogenic mechanisms specific to ADE compared to conventional infection suggests the opportunity for novel therapeutics leveraging strategies that manipulate host factors critical to infection by ADE. Thus, improving our understanding of mechanisms of ADE-mediated pathogenesis may provide insight for treatment approaches, vaccine development, and severe disease correlates for prognostication.

ADE has been demonstrated to result in elevated inflammatory pathology and increased virion production on a per cell basis using a variety of *in vitro* cell culture systems^14–17^. The elevated pathology observed in these models is thought to be in part mediated by increased efficiency of the FcγR interaction with DENV/IgG immune complexes relative to the interaction of DENV alone with canonical entry receptors such as DC-SIGN and the mannose receptor^18^. This high-affinity receptor interaction is thought to result in increased burden of infection due to increased virion entry to susceptible cells and has been recognized as hallmark “cell-extrinsic” features of ADE. The cell extrinsic effects of ADE are most often associated with *in vivo* DENV infections, where more and different cells are thought to be infected in the presence of enhancing antibody. In contrast, “cell intrinsic” response to infection by ADE has historically referred to altered signalling and cellular responses to infection which facilitate viral replication in ADE-infected cells^19^. Prior work has identified discrete transcriptomic and inflammatory signatures of intrinsic ADE, including the absence of an early interferon response and altered inflammatory signaling in cells infected by ADE ^15,17,20–22^. While the precise mechanisms underpinning these altered responses to infection remain incompletely understood, prior work points to preferential expression of anti-inflammatory cytokines (such as IL-10) and evasion of innate immune sensing (RIG-I, TLR)^15,23^. However, the existence of a cell-intrinsic ADE response – and our understanding of the transcriptomic mechanisms governing this response – has been a point of contention in the literature. The use of many different cell models and a variety of techniques for characterizing heterogeneous cultures has led to discordance in the proposed intrinsic mechanisms of ADE. It remains unclear whether the effects of enhanced infection occur primarily due to increased viral entry and exit, and how infection-elicited transcriptional profiles differ by route of viral entry.

Studies seeking to differentiate the cell-extrinsic from cell-intrinsic response to ADE during DENV infection have largely been impacted by technical limitations. However, recent advances in single cell RNA sequencing (scRNAseq) technology which allow for the capture of viral RNA in addition to host-derived transcripts have afforded the opportunity to identify and characterize features of infected cells in heterogeneous samples^24–27^. In this work, we applied a scRNAseq approach to specifically characterize the spectrum of cell-intrinsic responses to DENV infection in human monocyte-derived macrophages cultured under enhancing (ADE) or non-enhancing (conventional infection) conditions. This work identifies discrete transcriptional features of cells infected under different conditions of viral entry, where infected cells in ADE conditions demonstrated downregulated cytokine and interferon responses compared to those infected by conventional receptor-mediated entry. Uninfected/bystander cells in ADE cultures contrastingly displayed enrichment of immune signaling and responses, including increased cytokine production and antigen presentation. Together, these data provide insight into unique responses of infected and bystander cells under enhancing conditions and their role in DENV pathogenesis.

## RESULTS

### scRNAseq identifies distinct cell populations according to condition and infection status

To characterize the transcriptional profiles of DENV infected and bystander cells in various infection conditions, macrophages were differentiated from primary human PBMC-derived monocytes and infected with DENV-2 (strain NGC) in enhancing (ADE) or non-enhancing conditions, along with an uninfected control sample (**Methods, Fig. 1A**). The 10x Genomics 5’ scRNAseq assay was used for single cell partitioning and library generation, and a concatenated DENV-2/human reference genome was used to quantify both host cell and viral RNA transcripts. Base clustering of transcriptomic data identified 3 distinct clusters across cells from all three experimental conditions (**Fig. 1B**). These clusters corresponded broadly to cells from the control condition (cluster 1), and two clusters (cluster 0 and 2) with cells from both the conventional infection (DENV) and ADE (DENV+IgG) conditions (**Fig. 1B**). We observed the presence of elevated DENV RNA expression in cluster 2, and a low level DENV RNA signal in all cells in infected cultures (**Fig. 1C, Fig. S1**). This is consistent with previous reports indicating the presence of a low level of ambient RNA in bystander cells within infected cultures^26,28^. Based on patterns of DENV RNA and host transcriptomic expression, cells within cluster 2 were designated as infected (**Fig. 1D**). Using these infection state designations, 12.8% of cells in the DENV condition fell within the infected cell cluster, compared to 49.6% infection in DENV+IgG (ADE) conditions (**Fig. 4E**, **Table 1**). Consistent with previous reports, we observed an approximate 8-fold higher abundance of positive sense DENV RNA than negative sense RNA (**Fig. 1E**). DENV-infected cells in ADE conditions had a modest, but statistically significant increase in mean DENV RNA content on a per cell basis compared to cells infected under non-enhancing conditions (**Table 1**, **Fig. 1E**). Both DENV positive (+) and negative (-) sense RNA transcripts were detected within the dataset and were positively correlated across all infected cells (r = 0.96) (**Fig. 1F**). These findings support that we were successful in the capture of DENV and host cell transcripts under both conventional and IgG-enhanced infection conditions, enabling the differentiation of cell-intrinsic from cell-extrinsic ADE effects.

**Figure 1.**
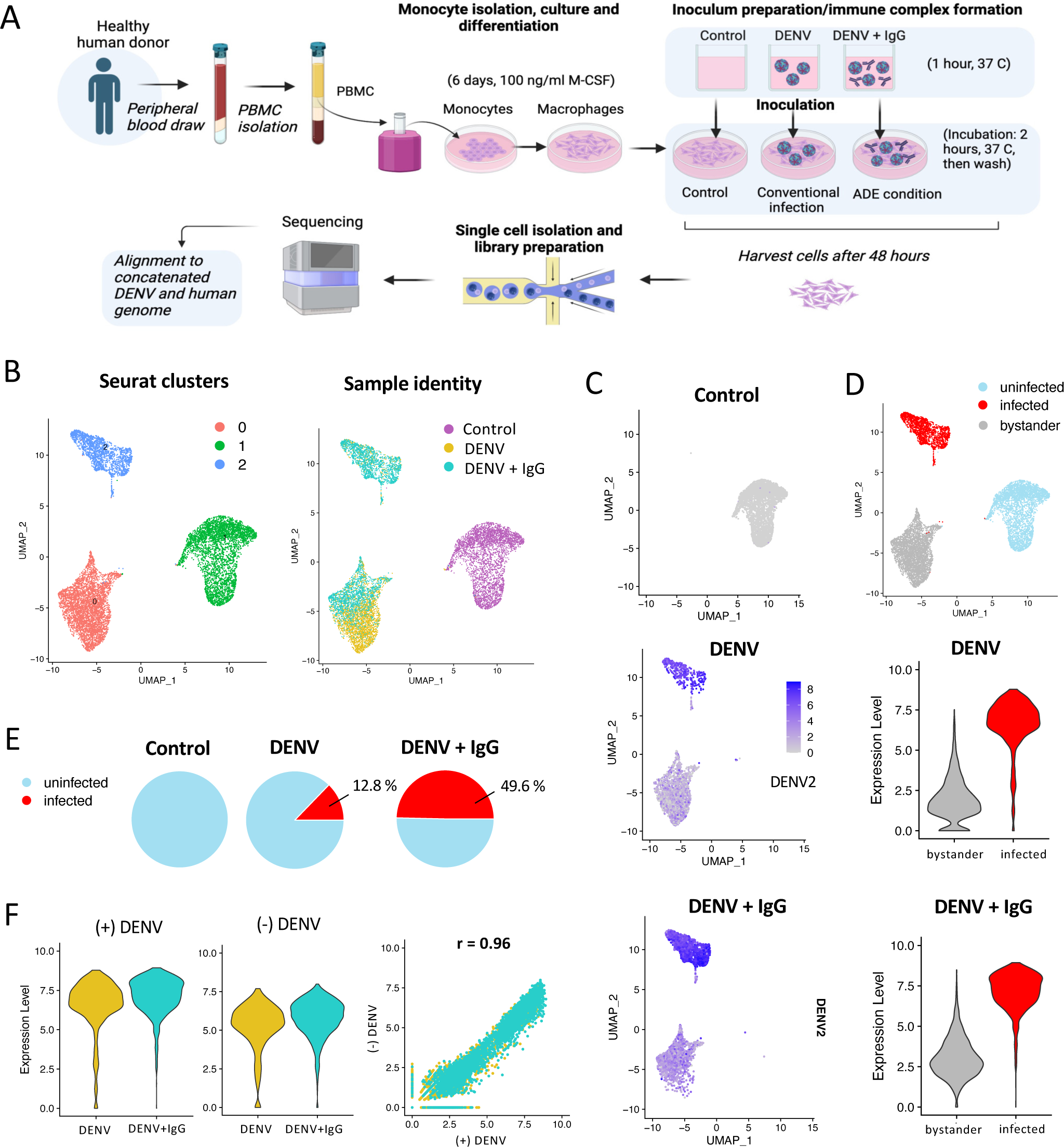
scRNAseq analysis of monocyte-derived macrophages in conventional infection or enhancing (ADE) conditions. **(A)** Schematic of experimental study design. Created in BioRender. **(B)** Integrated UMAP projections of scRNAseq data from all three experimental conditions: Control, DENV infected (conventional infection), and DENV+IgG (ADE condition) labelled according to Seurat clusters (clusters 0-3), sample origin (control, conventional infection, ADE). **(C)** Feature plot indicating DENV2 positive sense RNA expression across infection conditions. (**D**) Imputed infection state labelling of clusters, and corresponding violin plots demonstrating DENV2 positive sense RNA expression in infected, compared to bystander/uninfected cells in infected cultures. **(E)** Pie charts of infected cell proportions in control, DENV and DENV+IgG conditions. **(F)** DENV positive (+) and negative (-) sense RNA expression in infected cells and correlation plot of DENV (+) and DENV (-) RNA expression in infected cells.

**Table 1.** Infected cell characteristics across culture conditions.

| Sample | Infected cells (n) | Uninfected cells (n) | Total cells (n) | Percentage infected cells (%) | Mean viral RNA content per cell |  |
| --- | --- | --- | --- | --- | --- | --- |
|  |  |  |  |  | (+) sense | (-) sense |
| Control | 0 | 3740 | 3740 | 0 | 0 | 0 |
| DENV | 401 | 2741 | 3142 | 12.8 | 1389.0 | 365.3 |
| DENV+IgG | 1475 | 1496 | 2971 | 49.6 | 1937.8 | 526.7 |

### Infected and bystander cells demonstrate distinct gene expression profiles

To initially assess the differences in the transcriptomic profiles of DENV-infected and uninfected/bystander cells within heterogeneous infection cultures, differential gene expression analysis was performed between DENV-infected or uninfected/bystander cells and control cells (**Fig. 2A**). This analysis identified a total of 906 upregulated and 908 downregulated differentially expressed genes (DEGs) in infected cells compared to control (**Table S1, S2**). In bystander cells compared to control, 1270 genes were upregulated, and 1479 downregulated (**Table S3, S4)**. A core set of 639 upregulated and 819 downregulated genes were observed to be differentially expressed by cells in DENV-inoculated cultures relative to the control culture, irrespective of infection designation (**Fig. 2B**, **Table S5).** However, a number of DEGs were selectively differentially expressed within either the infected cells (352 genes) or uninfected/bystander cells (1291 genes) relative to control (**Fig. 2B, C, Table S6, S7**). Many of these infection-associated DEGs – either within the DENV-infected cells or the uninfected bystander cells within the same culture – correspond to canonical inflammatory/antiviral gene products, including cytokines, chemokines, immune signalling receptors and interferon stimulated genes (ISGs) (**Fig. 1C**).

**Figure 2.**
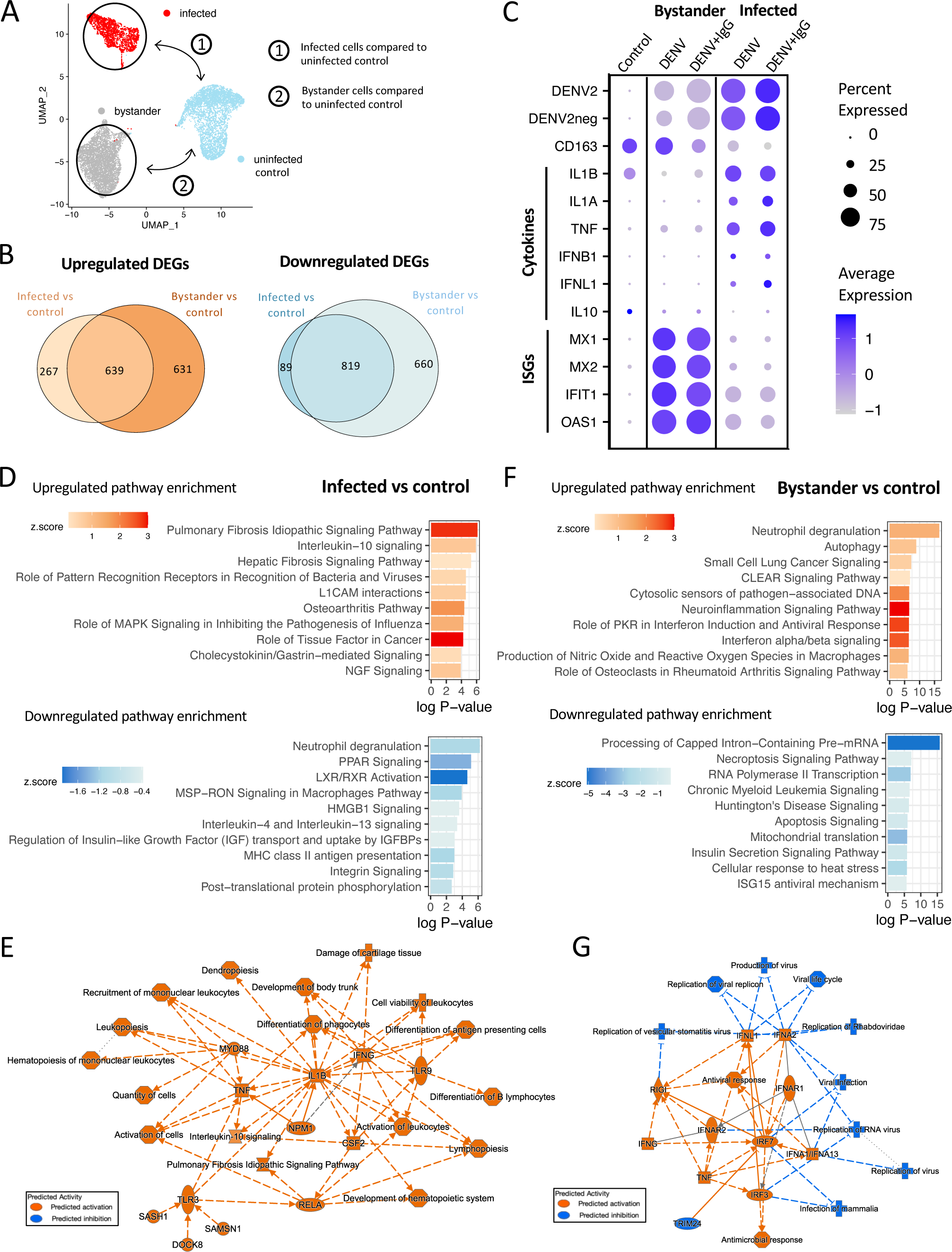
Transcriptomic profiles of infected and bystander cells. **(A)** UMAP of imputed labelling of cell populations indicating schematic of the cell populations being compared: infected cells compared to control, and bystander cells compared to control in DEG and IPA analysis **(B)** Euler plot depicting the number of differentially expressed genes (DEGs) in infected and bystander cells compared to control, values within plots represent the number of shared and unique DEGs in each comparison. **(C)** Dot plot indicating average expression of conventional infection markers, including various cytokines and interferon stimulated genes across conditions and infection states. Average expression indicated by colour intensity and percent of cells expressing a given transcript indicated by dot size. (**D**) Ingenuity Pathway Analysis (IPA) based on unique DEGs in Infected cells compared to control indicating top 10 upregulated and downregulated canonical pathways, and **(E)** graphical summary of IPA analysis in infected cells. **(F)** IPA based on unique DEGs in Bystander cells compared to control indicating top 10 upregulated and downregulated canonical pathways, and **(G)** graphical summary of IPA analysis in bystander cells. In graphical summary plots, red colour indicates activation, while blue indicates suppression. Solid lines represent direct interactions, dashed lines represent indirect relationships, and dotted lines represent inferred relationships.

DENV infected cells within the heterogeneous culture demonstrated selective upregulated expression of cytokines and chemokines such as *IL1A, IL1B, IFNB1,* and *TNF, IL6, w*hereas bystander cells demonstrated upregulation of ISGs such as *MX1, MX2, IFIT1, OAS*1 (**Fig. 1C**). Overall, these transcriptomic changes are consistent with expected signaling relationships, indicating activated pro-inflammatory signaling and interferon expression in infected cells, and response to these inflammatory cues by bystander uninfected cells in culture.

In order to determine associations of these gene expression changes as they relate to specific biological processes within DENV infected and bystander macrophages in DENV-inoculated cultures, we performed a pathway enrichment analysis using Ingenuity Pathway Analysis (IPA). Upregulated pathways in infected cells included elevated interleukin-10 signaling, pattern recognition receptor responses, growth factor signaling, and signaling associated with pathogenic processes (pulmonary fibrosis, hepatic fibrosis, osteoarthritis, cancer) (**Fig. 2D, Table S8**). IPA graphical summary analysis summarized the overall phenotype of infected cells as promoting proliferation and differentiation of lymphocytes and antigen presenting cells, immune cell recruitment, and differentiation of phagocytes **(Fig. 2E)**. The graphical summary of infected pathways interestingly emphasized the predicted role of TLR signaling and activation of IL-1β in driving these responses. Together, this analysis suggests an activated transcriptional profile specific to infected cells activated by innate immune receptors which promotes cytokine secretion, type I interferon induction, and proliferation, recruitment, and growth of leukocytes.

In uninfected/bystander cells compared to control, IPA analysis predicted many differentially regulated pathways, where similar to in DEG analysis, both the number of pathways and magnitude of these responses indicate a more prominent response in bystander than infected cells (**Fig. 2F, Table S9**). Of these pathways, top upregulated processes included activation of neutrophil degranulation, interferon signaling, and multiple pathways indicating activation of inflammation and anti-viral responses **(Fig. 2F)**. This was emphasized in graphical summary, where type I IFN (IFNA1, IFNA2, IFNA13) activation of IFNAR1/2 was predicted to induce anti-viral and ant-microbial responses **(Fig. 2G)**. These data also identified a role for IRF3 and IRF7-mediated activation of IFN signaling in bystander cells. Altogether, these findings support that the identified infected and uninfected/bystander cell populations represent transcriptionally distinct groups with expected immunological features according to infection state.

### Inflammatory signaling is decreased in macrophages infected in ADE conditions

Having defined the transcriptional profiles associated with DENV infection irrespective of route of infection, we next wanted to determine if there was any difference in DENV infection-elicited transcriptional profile between cells infected via ADE or conventional entry routes. Accordingly, we performed differential gene expression analysis specifically on cells within the DENV infected cluster, binning by infection condition (DENV+IgG vs DENV only) (**Fig. 3A**). This analysis identified a total of 162 differentially expressed genes within the infected cell cluster between the two infection conditions, of which 45 genes were significantly upregulated, and 117 genes were significantly downregulated (**Table S10, S11**). Top downregulated DEGs included many chemokine ligands such as *CCL8*, *CCL2*, *CXCL10*, *CCL7*, and *CXCL11*, and immune signaling associated genes such as *P2RX7, STAT1*, and *JAK2* **(Fig. 3B, Table S11)**. Although there appeared to be overall immunosuppressive changes to gene expression in cells infected by ADE compared to conventional infection, select immune signalling molecules were enriched, including top upregulated DEGs *CCR7*, *ISG15*, *IL23A*, and *TRAF* (**Fig. 3B**).

**Figure 3.**
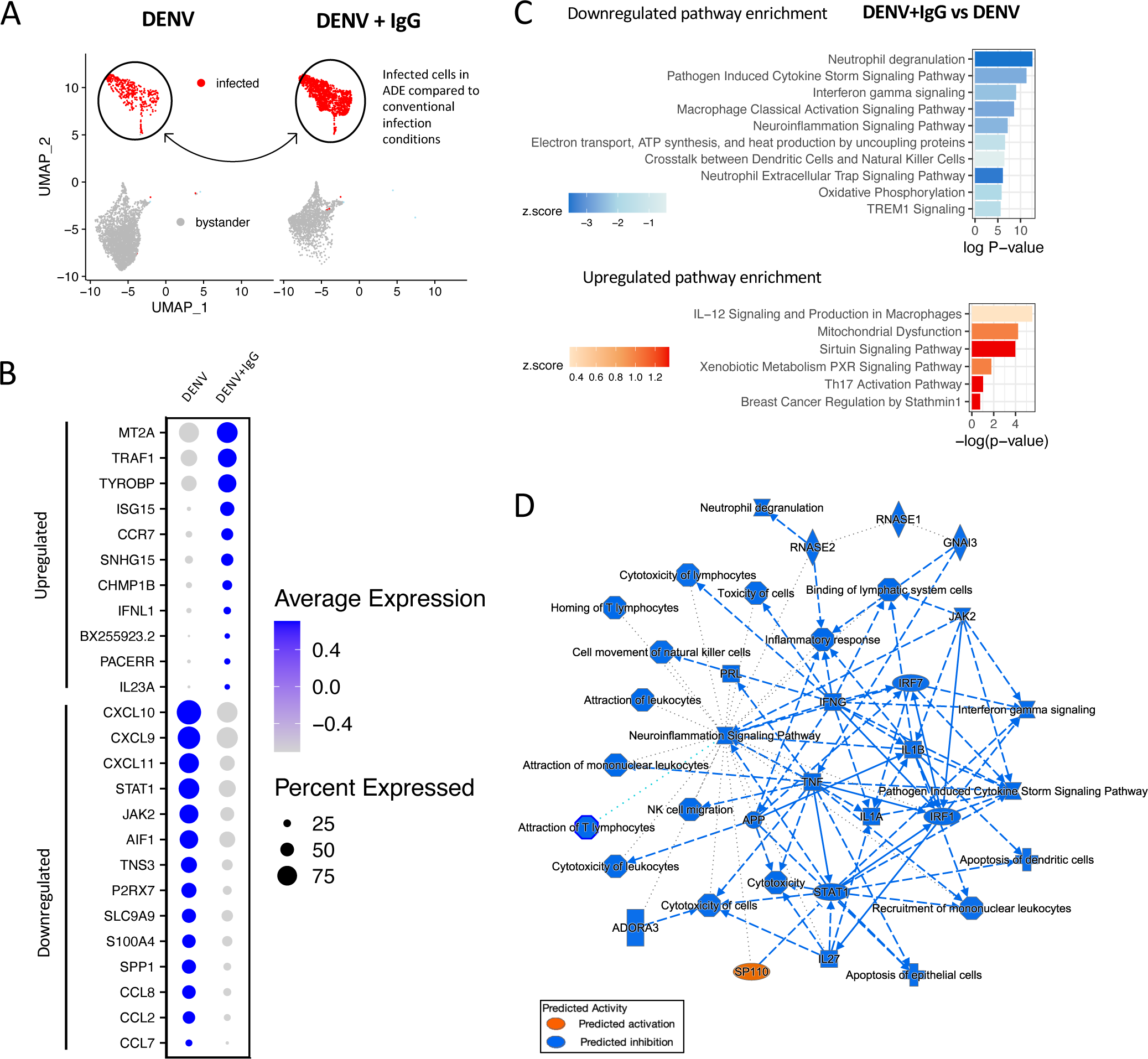
Gene expression and ingenuity pathway analysis of infected macrophages in ADE compared to conventional infection conditions. **(A)** Schematic of comparison being made between infected cells in DENV+IgG (ADE) compared to DENV (conventional infection) conditions. **(B)** Dot plot highlighting selectively upregulated and downregulated DEGs in ADE compared to conventional infection, indicating average expression and percent of cells expressing a given transcript. (**C)** IPA based on DEGs in infected cells in ADE compared to conventional infection. The 6 upregulated and top 10 downregulated canonical pathways are displayed. **(D)** Graphical summary of IPA analysis in infected cells. In graphical summary plots, red colour indicates activation, while blue indicates suppression. Solid lines represent direct interactions, dashed lines represent indirect relationships, and dotted lines represent inferred relationships.

Pathway analysis was performed using an input of DEGs between infected cells in ADE compared to conventional DENV infection conditions. In infected cells, IPA analysis predicted a predominantly suppressed phenotype in ADE compared to conventionally infected cells (**Fig. 3C, Table S12**). Pathway analysis notably indicated downregulation of the inflammatory response in ADE-infected cells, where downregulated pathways included: neutrophil degranulation, pathogen induced cytokine storm signaling pathway, interferon gamma signaling, and macrophage classical activation signalling pathway (**Fig. 3C**). Pathways involved in leukocyte recruitment, activation, and proliferation were downregulated compared to cells infected in conventional infection conditions. There also appeared to be an enriched signature of mitochondrial dysfunction in ADE-infected cells, and downregulated pathways including electron transport, ATP synthesis and heat production by uncoupling proteins, and oxidative phosphorylation (**Fig. 3C**). Of the few upregulated pathways in cells infected in ADE conditions, IL-12 signaling interestingly emerged as a top pathway. In graphical summary analysis these processes were summarized as being driven by decreases in cytokine production such as IL-1α, IL-1β, and TNF, and decreased interferon response signaling by STAT1 and JAK2 (**Fig. 3D**).

These reflect both a decrease in innate anti-viral activation of cytokine responses as well as secondary immune response to IFN production. Together these processes were highlighted as contributing to downregulated recruitment, activation, and cytotoxicity of leukocytes, neutrophil degranulation, IFN-γ and cytokine storm signalling (**Fig. 3D**). The observed decrease in pro-inflammatory processes in macrophages infected in ADE conditions might indicate mechanisms of attenuated immune response in cells infected by this route of entry which facilitate viral replication and contribute to enhanced viral burden.

### ADE-associated gene expression changes are independent of DENV RNA content

Given that infected cells in ADE conditions demonstrated increased mean viral RNA on a per cell basis, we next sought to determine whether the transcriptomic changes observed in DENV infected cells under ADE conditions were simply due to increased viral burden within cells. To this end, correlation analysis was performed between DENV RNA content and expression of all genes in the dataset within the infected cells. Across all host genes captured in this analysis, no significant correlation between DENV (+) RNA expression and the expression of cellular transcripts was observed (**Fig. 4A**). This observation was also true for the expression of DENV (-) RNA and cellular transcripts. Specific interrogation of top DEGs between infected cells from ADE compared to conventional infection conditions also demonstrated no significant correlation with viral RNA content in infected cells (**Fig. 4B-D**). These findings indicate that transcriptomic differences are likely not driven by differences in DENV RNA content and support that these changes are instead reflective of intrinsic cell differences in ADE and conventional infection.

**Figure 4.**
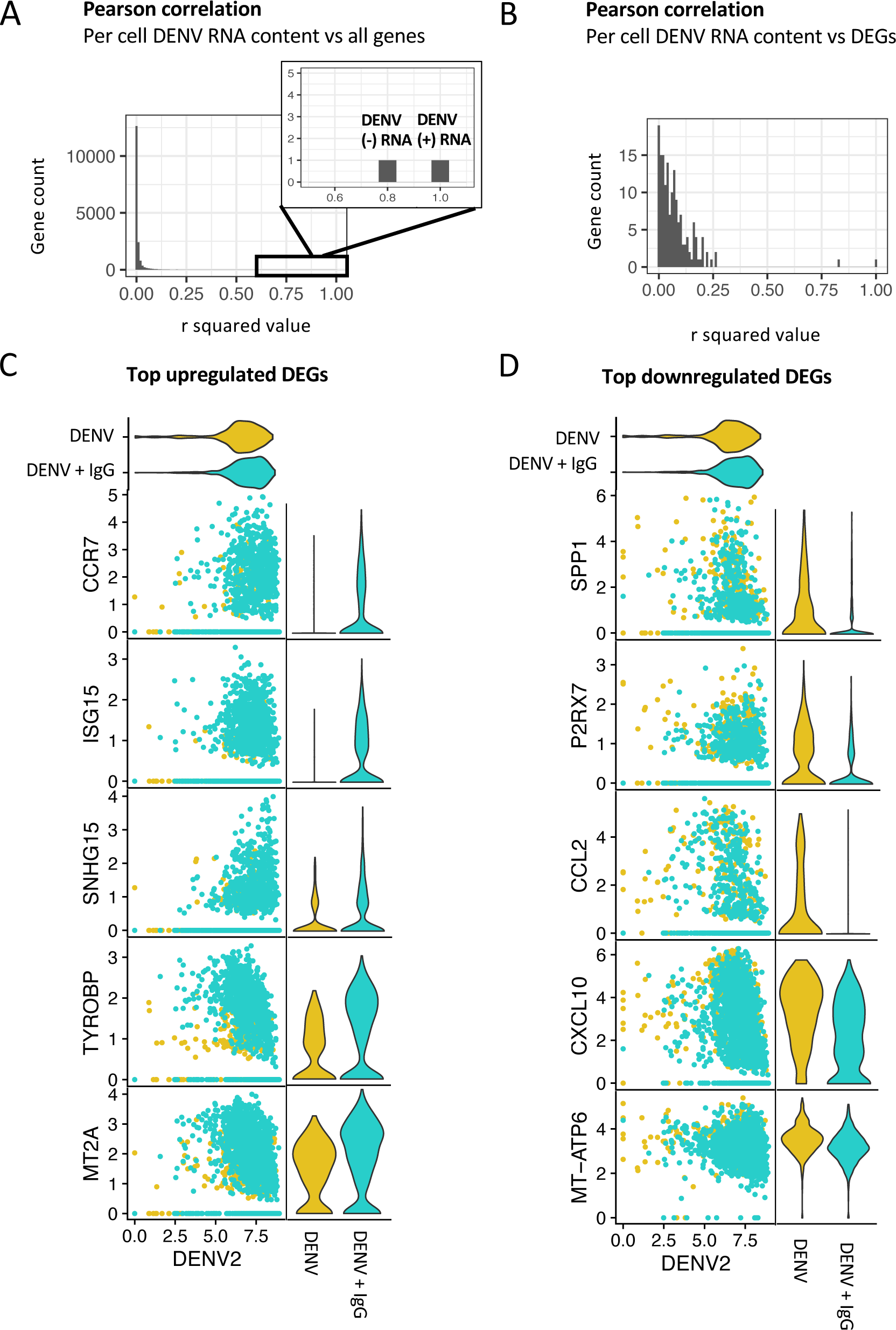
Correlation of DENV RNA expression with gene expression in infected cells. **(A)** Histogram with bins indicating the number of genes with a given r-squared value in Pearson correlation analysis of DENV RNA expression with each gene in the dataset, specifically in infected cells. Right hand upper corner of the plot is zoomed in to indicate R-squared values for DENV (+) and (–) sense RNA. **(B)** Histogram with bins indicating the number of genes with a given r-squared value in Pearson correlation analysis of DENV RNA expression with that of DEGs differentially expressed in infected cells from ADE compared to conventional infection conditions. **(C)** Pearson correlation of DENV (+) RNA expression with expression of select top upregulated DEGs from comparison of infected cells in ADE compared to conventional infection conditions. Corresponding violin plots of upregulated DEG expression in DENV (yellow) and DENV+IgG (turquoise) conditions. **(D)** Pearson correlation of DENV (+) RNA expression with expression of select top downregulated DEGs from comparison of infected cells in ADE compared to conventional infection conditions. Corresponding violin plots of downregulated DEG expression in DENV (yellow) and DENV+IgG (turquoise) conditions.

### Bystander cells in ADE conditions have heightened anti-viral responses

Although most of the work defining intrinsic ADE has focused on characterizing either bulk cellular responses or those of infected cells, the potentially unique features of bystander cells in ADE conditions have yet to be explored. We identified discrete differences in the responses of infected and bystander cells across conditions, and therefore postulated that bystander cells in different infection conditions also possessed unique transcriptomic features. We next looked specifically at bystander cells as designated by initial clustering and imputed cell labelling to evaluate gene expression differences between cells in ADE (DENV+IgG) and conventional infection (DENV) conditions (**Fig. 5A**). In this comparison, DEG analysis identified 467 differentially expressed genes, of which 238 were upregulated and 229 were downregulated (**Table S13, 14).** Notably, these changes along with those identified in pathway analysis indicated that bystander cells harbored more distinct changes between infection conditions than infected cells. Among top upregulated DEGs were *TYROBP, ISG15, IFITM3, RPL28, CD74, CCR7* **(Fig. 5B**). Interestingly, *TYROBP, ISG15* and *CCR7* were DEGs in common between infected and bystander cells in ADE compared to conventional infection conditions. Bystander cells in ADE compared to conventional infection conditions uniquely demonstrated significant upregulation of MHC expression. Top downregulated DEGs in bystander cells indicated a decreased enrichment of certain pro-inflammatory cytokines and signaling molecules in ADE compared to conventional infection conditions, including *CXCL11, CXCL10, CCL2, DOCK4, STAT1*, and *JAK2* **(Fig. 5B**). Notably, while many gene expression changes were unique to infected or bystander cells, a number of up or downregulated genes were common between infected and bystander cells in ADE compared to conventional infection conditions. For example, *TYROBP, CCR7, CXCL10,* and *CXCL11* emerged as top DEGs in both infected and bystander cells in ADE compared to conventional infection.

**Figure 5.**
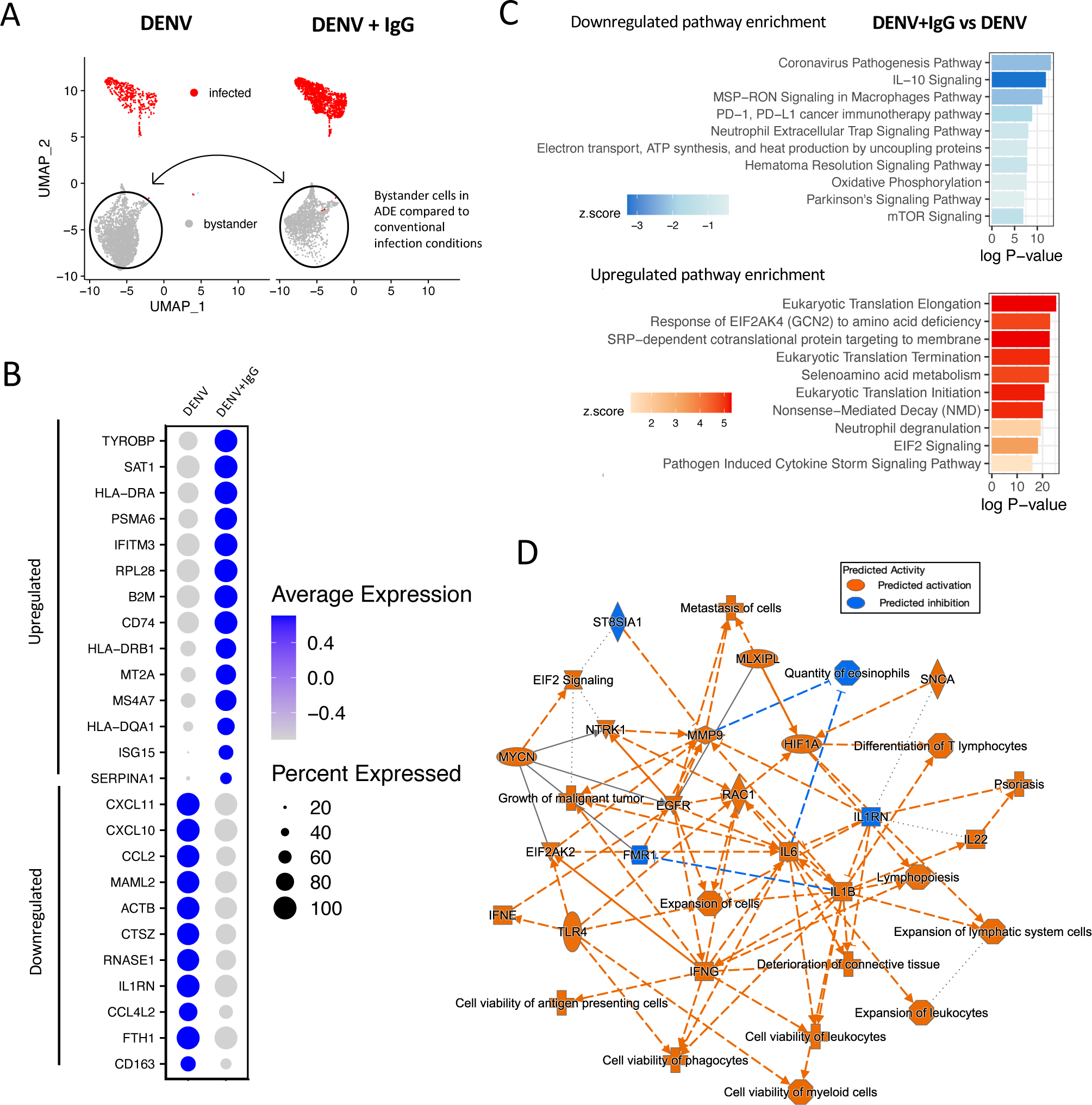
Gene expression and ingenuity pathway analysis of bystander cells within infected cultures in ADE compared to conventional infection conditions. **(A)** Schematic of comparison being made between bystander cells in DENV+IgG (ADE) compared to DENV (conventional infection) conditions. **(B)** Dot plot highlighting selectively upregulated and downregulated DEGs in ADE compared to conventional infection, indicating average expression and percent of cells expressing a given transcript. (**C)** IPA based on DEGs in bystander cells in ADE compared to conventional infection conditions. Top 10 upregulated and downregulated canonical pathways displayed. **(D)** graphical summary of IPA analysis in bystander cells. In graphical summary plots, red colour indicates activation, while blue indicates suppression. Solid lines represent direct interactions, dashed lines represent indirect relationships, and dotted lines represent inferred relationships.

In bystander cells, IPA analysis identified enriched pathways including those associated with neutrophil degranulation and cytokine storm signaling, in addition to a number of pathways relating to translation initiation and elongation **(Fig. 5C**). Pathway analysis also highlighted a role for antigen presentation pathway upregulation, consistent with the MHC gene expression changes described (**Table S15**). Downregulated IL-10 signaling emerged as a top downregulated pathway in bystander cells in ADE conditions, and similar to in infected cells, a prominent role for downregulation of metabolic processes involving electron transport and oxidative phosphorylation **(Fig. 5C**). Integration of pathway enrichment data in graphical summary demonstrated increased cell viability of phagocytes and APCs, differentiation of lymphocytes, and EIF2 signaling in bystander cells in ADE conditions. These changes were driven by increases in transcript production of various signaling molecules such as *HIF1A, EGFR, IL22, IFNE, IFNG, TLR4, IL6,* and *IL1B* (**Fig. 5D**). Compared to bystander cells in conventional infection conditions, those in ADE conditions demonstrated an overall activating response by these processes. These findings indicate that in ADE conditions alternative signaling by infected cells or increased infection burden contribute to distinct differences in bystander cell responses which may provide insight to our understanding of cellular responses to DENV infection.

## DISCUSSION

In this study, we applied a scRNAseq approach to determine the transcriptomic profiles of infected cells in ADE and conventional infection conditions in order to characterize the cellular response to differing routes of DENV infection. We found that cells infected by ADE compared to conventional infection demonstrated a discernable transcriptomic profile characterized by decreased immune signalling, IFN response, and mitochondrial dysfunction. Further, we demonstrate that the observed gene expression and pathway analysis signatures are independent of DENV RNA content, indicating that the observed changes are not simply due to increased viral burden per cell in ADE conditions. We also identify distinct profiles in bystander cells in ADE compared to conventional infection conditions, indicating an activated phenotype with enriched protein translation, cytokine production, and antigen presentation. This work provides novel insight into the cell intrinsic mechanisms, specifically in infected and bystander cells, which may be responsible for enhanced infection in ADE. The unique profiles of bystander and infected cells within and across conditions highlights the utility and importance of dissecting these profiles in a heterogenous cell population.

The dynamic interplay of virus and host responses is demonstrated by the complex signaling and variable responses of infected and bystander cells to infection. These unique features of infected and bystander cells have been elegantly described with single cell resolution in a number of pathogens such as influenza, Zika virus (ZIKV), and West Nile Virus (WNV)^25,29–33^. Our study characterizes the comprehensive transcriptomic profiles of infected and bystander human macrophages in response to DENV infection, contributing to the body of literature describing these divergent responses. Specifically in the context of intrinsic ADE, most studies investigating cell intrinsic mechanisms have done so in bulk cultures or have focused on characterizing only the profiles of the infected cells. To our knowledge, we report the first unique characterization of the features of bystander/uninfected cells in ADE conditions. This distinction further lends insights into how bystander cell profiles may contribute to and bias findings when looking at the level of whole cultures.

Work by Hamlin et al. in human dendritic cells described an increase in TNF and IL-1β production from DENV infected compared to bystander cells, which upregulate the expression of ISG IP-10^29^. We report similar findings in this single cell analysis of DENV infection of macrophages, where bystander cells took on an anti-viral state, as has been described in other studies in other pathogens^26,29,30,34,35^. Hamlin et al. comment that prior studies characterizing infected cell responses by bulk culture analysis indicating cell adoption of an anti-viral state are likely emanating the profiles of the bystander, rather than infected cells in culture^29^. Given the increased magnitude of response that we observe in bystander compared to infected cells and the dominant activating phenotype of these cells, our data also support this observation. We saw IFN-β to be a uniquely upregulated gene in infected cells, consistent with recent findings of Moore et al. in scRNAseq analysis of Zika virus (ZIKV) infection, where they report that IFN-β expression as a unique feature of infected cells and driving force for the inflammatory pathology in ZIKV infection. Moore et al. identify a strong correlation between ZIKV RNA content and *IFNB* gene expression, however, we observed no such relationship between DENV RNA content and expression of any genes in our dataset^26^. These differences might be due to differences in timing of infection or indicate unique features of IFN response in these two pathogens.

When comparing the gene expression profiles of infected or bystander cells between enhancing and non-enhancing conditions we identified a number of differentially regulated processes in cells infected in ADE conditions (**Fig. 6)**. Previous work has highlighted a potential role for decreased type I and II IFN and proinflammatory cytokine responses in ADE-mediated infection^11,15,22^. Some of this work has very specifically focused on a role for attenuated IFN-β production in cells infected by ADE. Although *IFNB* did not explicitly emerge as a key DEG between infected cells in ADE compared to conventional infection conditions, we observe a trend toward decreased *IFNB* transcript production. Similar to previous reports, we identify an overarching signature indicating suppression of immune responses in cells infected by ADE. Our data indicates a likely role for differential cellular response to IFN responses, rather than abrogating innate immune recognition and initial type I IFN induction. It is known that in conventional infection, DENV2 evades innate immune activation and IFN responses by a number of mechanisms, including its non-structural protein NS3’s inhibition of STAT1 and STAT2 signaling^36–38^. Given that STAT1 and JAK2 were significantly downregulated in ADE compared to conventionally infected cells, it is possible that in ADE-mediated infection these interactions are differentially modulated such that they further enhance viral evasion strategies in ADE. Increased IL-10 production has also been highly proposed as a mechanism of silencing immune activation in ADE conditions or individuals with severe dengue^15,20,23^. Interestingly, our data demonstrated no significant elevation of IL-10 transcript production in infected or bystander cells in ADE conditions. In fact, in IPA analysis IL-10 signaling emerged as one of the top downregulated pathways in bystander cells in ADE conditions compared to those in conventional infection.

**Figure 6.**
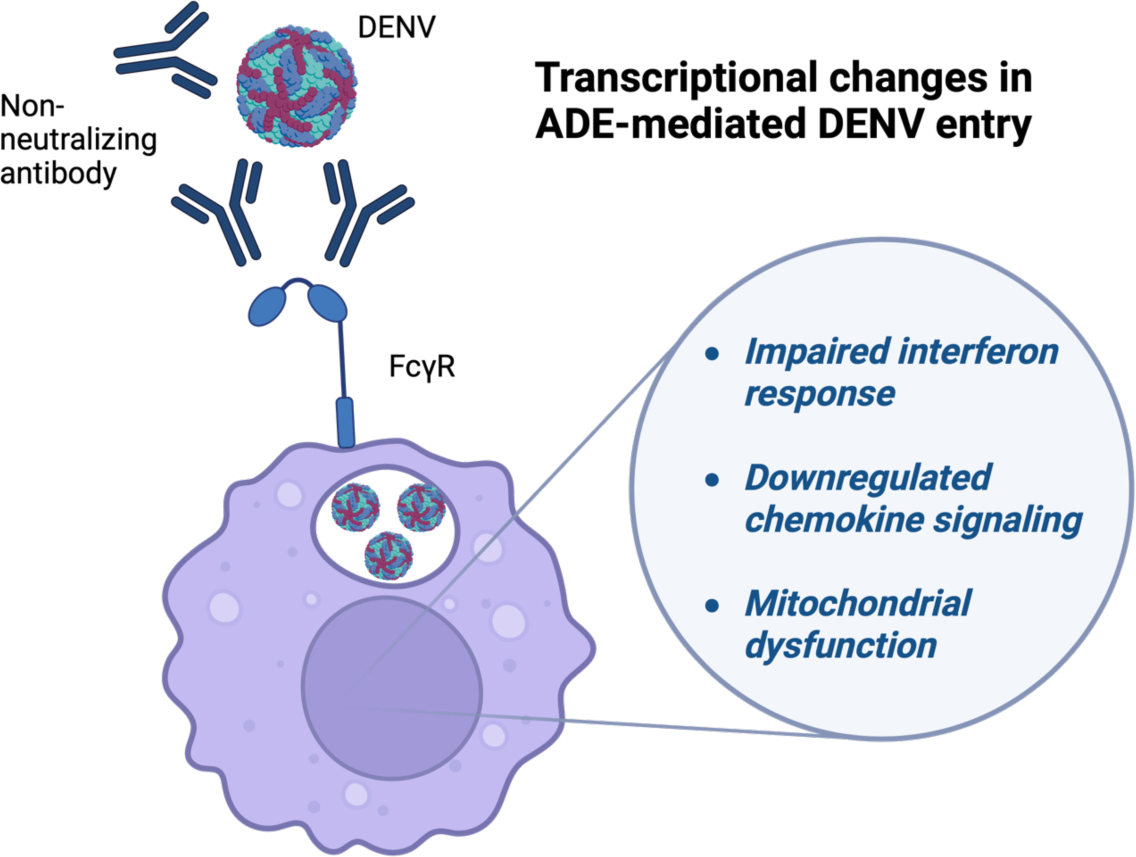
Summary of transcriptomic features of infected cells in ADE conditions. Human macrophages infected in the presence of enhancing antibodies demonstrate distinct gene expression profiles indicating mitochondrial dysfunction, suppressed interferon responses and decreased inflammatory chemokine transcription compared to those in conventional infection conditions. Created in BioRender.

One of the most recent studies assessing a role for intrinsic ADE identified elevated protein translation and splicing as an intrinsic feature of cells infected by ADE^39^. Although we did not observe this same finding in infected cells, elevated protein translation was observed as a feature of bystander cells in ADE conditions. Our data also suggest a prominent role for dysfunctional mitochondrial responses in cells infected in enhancing conditions (**Fig. 3C**). Interestingly, these findings are consistent with observations by Chan et al., who also note trends in upregulation of mitochondrial respiratory chain complexes specifically in cells infected by ADE^39^. Our analysis also indicated novel specific downregulation of chemokines which have not specifically been reported. IPA analysis summarizes these effects as driving an overall decrease in immune cell homing and recruitment, reducing natural killer cell, lymphocyte, and neutrophil migration and killing. To our knowledge, prior mechanisms have not directly linked or suggested altered chemokine responses and cellular migration to be specific hallmark effects of ADE. Altogether, our analysis fortifies a number of previously suggested mechanisms of intrinsic ADE in addition to suggesting novel changes in gene expression unique to cells in ADE conditions.

It has remained difficult to define which proportion of differences in response are due to alternative receptor engagement, differences in virus compartmentalization by route of entry, or simply driven by differences in viral load in culture. Discordance in the literature regarding mechanisms of intrinsic ADE may be due to the fact that it is impossible to truly distinguish intrinsic from extrinsic features of enhanced infection in realistic physiologic settings. While various methodologies have been employed to control for differences in viral burden, these impact one of the fundamental features of antibody mediated entry which is that it elevates virion production and inflammatory pathology. In our analysis we did not compensate for elevated infection in ADE conditions, because we aimed to evaluate together the intrinsic and extrinsic effects of ADE. We provide a comprehensive analysis of the transcriptional responses of DENV-infected macrophages; however, we recognize that it is possible that alternatively modulated processes might be post-transcriptionally mediated, changes which would not be captured in our analysis. IPA graphical summary analysis allowed a glimpse into predicted relationships and alterations based on pathway alterations at the transcriptomic level, but additional insight is necessary to understand the mechanisms at the post transcriptional, and protein level. Finally, we recognize that these findings were confined to a single DENV serotype and strain, and at a single time point, in an isolated in vitro system that does not allow the assessment of the impact of various interactions with other immune cells.

Future work might seek to understand the functional impact of the observed changes in a less isolated system, determining the effects of different cell phenotypes in other leukocytes, B cells and T cells. It would be meaningful for additional studies to assess whether similar signals can be identified in natural infection or human infection model samples. ADE is a feature of viruses such as ZIKV and SARS-CoV2, where application of similar methodology would provide useful insight into the underlying intrinsic mechanism, how, and whether these differ in ADE of other pathogens ^40–42^. In this study, we describe evidence of novel pathways and mechanisms which may be driving enhancement in both DENV infected and bystander cells in ADE infection conditions. This work provides the groundwork for additional study of these mechanisms, their contribution to ADE pathogenesis, and potential as targets for interventional approaches.

## METHODS

### Viruses

DENV-2 (strain New Guinea C) stocks were prepared by serial passaging in Vero cells. Plaque assay of Vero cells was used to determine stock infectious titer (PFU) for each virus. An MOI of 1 was used in all infection assays.

### Monocyte-derived macrophage differentiation

Peripheral blood mononuclear cells (PBMC) were isolated and cryopreserved from whole blood of healthy human donors. Monocytes were isolated from PBMC and differentiated into macrophages as previously described by Wegman et al ^20,43^. The Mojosort monocyte isolation kit was used to isolate monocytes by negative selection according to the manufacture’s protocol (Biolegend, #480059) and cells were resuspended in RPMI supplemented to with 100ng/mL M-CSF (Peprotech, # 300-25), 10% (v/v) FBS (Gibco), 1% penicillin/streptomycin, 1% L-glutamine. Cells were plated on tissue culture treated 24 well plates at a cell density of 1.25-1.5×10^6^ cells per mL. In order to ensure monocyte adhesion, plates were briefly centrifuged for 2 minutes at 500 g. Day 1 was defined as the day of isolation and plating. Cells were subsequently incubated at 37°C/5% CO_2_ with culture media repletion on day 5 by adding 1 mL of differentiation media per well.

### DENV infection of monocyte-derived macrophages

In order to allow immune complex formation, inocula were prepared by incubating DENV-2 with VDB33 monoclonal IgG (ADE condition), DENV-2 alone with media (conventional infection condition), or media alone (uninfected control) at 37 °C/5% CO_2_ for 60 minutes. An antibody concentration of 1 ug/mL, previously demonstrated to induce enhancing conditions in primary human macrophages, was used^43,44^. On day 6 of culture, supernatants were aspirated from monocyte-derived macrophage cultures and 200 uL of inoculum was added to each well according to desired infection conditions. Cells were incubated with inoculum for 2 hours at 37°C/5% CO_2_, inoculum was removed, and cells were washed twice with 2mL of RPMI medium. 500 mL of differentiation media was added to each well and plates incubated for 48 hours at 37C/5% CO_2_. Macrophages were then detached from plates by adding 600 mL of Accutase (Stemcell technologies, 07920) per well and incubated at 37°C for 20 minutes. Cells were dissociated by pipetting and transferred to15 mL conical tubes and analyzed as described.

### Single cell RNA sequencing

Samples were prepared for single-cell RNA sequencing according to the 10x Genomics RNA-seq protocol. Cells were resuspended at a concentration of 500 cells/uL in PBS and loaded for a target of 6000 cells per reaction. Cells were loaded for Gel emulsion bead (GEM) generation and barcoding. Construction of 5’ gene expression libraries was performed using the Next GEM Single Cell 5’ reagent kit, Library Construction Kit, and The i7 Multiplex Kit (10x Genomics, CA). was used for reverse transcription, complementary DNA amplification and construction of gene expression libraries. The quality of gene expression libraries was assessed using an Agilent 4200 TapeStation with High Sensitivity D500 ScreenTape Assay and Qubit Fluorometer (Thermo Fisher Scientific) according to the manufacturer’s recommendations. Sequencing of 5’ gene expression libraries was performed on an Illumina NextSeq 2000 (Illumina) using P3 reagent kits (100 cycles). Parameters for sequencing were set at 26 cycles for Read1, 8 cycles for Index1, and 98 cycles for Read2.

### 5’ gene expression analysis and visualization

The 10x Genomics Cell Ranger pipeline was used to perform gene expression alignment. Sample demultiplexing, alignment, and barcode/UMI filtering was performed using the Cell Ranger software package (10x Genomics, CA) and bcl2fastq (Illumina, CA) using the commands mkfastq and count. A reference genome was created by combining the human reference genome (Ensembl GRCh38.93) with the DENV2 genome as an additional chromosome (NC 001474.2). Sequenced transcripts were aligned to a human reference library created using the Cell Ranger mkref command, combined human and DENV2 reference genome, and custom Ensembl GRCh38 DENV2 GTF. Reads were mapped to both the positive and negative sense DENV2 genome.

Multi-sample integration, data normalization, dimensional reduction, visualization and differential gene expression were performed in R studio (v4.3.2) using R package Seurat (v4.4.0). Sample datasets were filtered to contain cells with less than 22% mitochondrial RNA content and between 400-3,500 unique features. Genes expressed in fewer than 3 cells were excluded from analyses. The resulting dataset was normalized and scaled using the Seurat functions NormalizeData(), ScaleData(), and FindVariableFeatures(). Following data normalization and scaling, principal component analysis was performed using RunPCA(). Cells were then clustered based on the first 10 principal components and with a resolution parameter of 0.1 using FindNeighbors() and FindClusters(), respectively. Initial clustering indicated the presence of five clusters, of which two very small, but distinct clusters were characterized by B and T cell marker gene expression. We expect that these clusters represent a small number of contaminating cells which remained following monocyte isolation from PBMC. These clusters were thus excluded from subsequent analysis.

### Statistical analysis

Differentially expressed genes were identified by applying the FindMarkers() command and the Wilcoxon rank-sum test to the normalized gene expression dataset. A default minimum logFC value of 0.1 and min.pct of 0.01 were used. Adjusted p-values based on Bonferroni correction using all genes in the dataset. Genes with a corrected p-value of <0.05 were considered significant. To determine mechanisms associated with gene expression changes, Ingenuity Pathway Analysis (IPA, Qiagen) was performed. Analysis was performed using genes with a corrected p-value <0.01 and log fold change <-0.25 or >0.25. IPA results with a p-value <0.01 and z-score <-0.25 or >0.25 were considered statistically significant.

## Supplemental material

**Figure S1.** DENV RNA content across infection conditions.

**Table S1.** Upregulated DEGs in infected cells compared to uninfected control. **Table S2.** Downregulated DEGs in infected cells compared to uninfected control. **Table S3.** Upregulated DEGs in bystander cells compared to uninfected control.

**Table S4.** Downregulated DEGs in bystander cells compared to uninfected control.

**Table S5.** Common DEGs in comparisons between infected compared to uninfected control, and bystander compared to uninfected control.

**Table S6.** DEGs unique to the infected vs uninfected control comparison.

**Table S7.** DEGs unique to the bystander vs uninfected control comparison.

**Table S8.** IPA analysis of differentially enriched pathways in infected cells compared to uninfected control.

**Table S9.** IPA analysis of differentially enriched pathways in bystander cells compared to uninfected control.

**Table S10.** Upregulated DEGs in infected cells in ADE compared to conventional infection conditions.

**Table S11.** Downregulated DEGs in infected cells in ADE compared to conventional infection conditions.

**Table S12.** IPA canonical pathways differentially enriched between infected cells in ADE compared to conventional infection conditions.

**Table S13.** Upregulated DEGs in bystander cells in ADE compared to conventional infection conditions.

**Table S14.** Downregulated DEGs in bystander cells in ADE compared to conventional infection conditions.

**Table S15.** IPA canonical pathways differentially enriched between bystander cells in ADE compared to conventional infection conditions.

## ACKNOWLEDGEMENTS

We gratefully acknowledge the technical assistance of Karen Gentile of the Upstate Medical University Molecular Analysis Core (MAC). We thank Lauren Bahr and Michael McCracken for their editing and review of the manuscript.

## Author contributions

Céline S. C. Hardy, Conceptualization, Data curation, Formal analysis, Investigation, Methodology, Visualization, Writing – original draft, Writing – review and editing | Adam D. Wegman, Methodology, Writing – review and editing | Mitchell Waldran, Methodology, Writing – review and editing | Adam T. Waickman, Conceptualization, Data curation, Formal analysis, Investigation, Methodology, Funding acquisition, Supervision, Writing – review and editing.

## Data availability

scRNAseq data are publicly available through the Gene Expression Omnibus. Code used to perform scRNAseq analysis in R is available upon request.

## Funding

Funding for this study was provided by the State of New York and the Military Infectious Disease Research Program.

